# A short course of Tofacitinib sustains the immunoregulatory effect of CTLA4-Ig in presence of inflammatory cytokines and promotes long-term survival of murine cardiac allografts

**DOI:** 10.1101/2020.11.18.388595

**Authors:** Marcos Iglesias, Saami Khalifian, Byoung Chol Oh, Yichuan Zhang, Devin Miller, Sarah Beck, Gerald Brandacher, Giorgio Raimondi

**Author notes:** Corresponding authors e-mail: Giorgio Raimondi and Marcos Iglesias. These authors contributed equally to this work.

## Abstract

Costimulation blockade-based regimens are a promising strategy for management of transplant recipients. However, maintenance immunosuppression via CTLA4-Ig monotherapy is characterized by high frequency of rejection episodes. Recent evidence suggests that inflammatory cytokines contribute to alloreactive T cell activation in a CD28-independent manner, a reasonable contributor to the limited efficacy of CTLA4-Ig. In this study, we investigated the possible synergism of a combined short-term inhibition of cytokine signaling and CD28 engagement on the modulation of rejection. Our results demonstrate that the JAK/STAT inhibitor Tofacitinib restored the immunomodulatory effect of CTLA4-Ig on mouse alloreactive T cells in presence of inflammatory cytokines. Tofacitinib exposure conferred dendritic cells with a tolerogenic phenotype reducing their cytokine secretion and costimulatory molecules expression. JAK inhibition also directly affected T cell activation. In vivo, the combination of CTLA4-Ig and Tofacitinib induced long-term survival of heart allografts and, importantly, it was equally effective when using grafts subjected to prolonged ischemia. Transplant survival correlated with a reduction in effector T cells and intragraft accumulation of regulatory T cells. Collectively, our studies demonstrate a powerful synergism between CTLA4-Ig and Tofacitinib and suggest their combined use is a promising strategy for improved management of transplanted patients.

## 1. Introduction

Immunosuppression minimization and withdrawal through tolerance induction remain the focus of transplant research for better management of transplanted patients.^1,2^ Costimulation blockade strategies are a promising approach to induce tolerance to allogeneic tissues.^3-7^ A prototypical example is cytotoxic T-lymphocyte associated protein 4 (CTLA4)-Ig, a biologic used to prevent the interaction of CD80 and CD86 on antigen presenting cells (APCs) with CD28 on T cells. The consequent T cell receptor (TCR) engagement (Signal 1), without CD28 costimulation (Signal 2), results in decreased IL-2 production and actuation of a program of apoptosis or induction of T cell anergy.^8^ However, CD28 blockade via CTLA4-Ig alone does not prevent all T cell proliferation and is not sufficient to promote tolerance.^9^ Indeed, multiple clinical studies have confirmed that CTLA4-Ig monotherapy is characterized by high frequency of rejection episodes.^2,10,11^

The combined blockade of CD28 and CD40 signaling pathways was originally proposed as the strategy to achieve lasting tolerance in multiple rodent models of transplantation.^9^ However, a clinical trial with a humanized antibody to CD154 (the ligand of CD40) was halted due to thromboembolic events.^12^ Interestingly, blockade of the CD40/CD154 axis inhibits APC secretion of inflammatory cytokines,^13^ and multiple studies have demonstrated that acute inflammatory events (e.g. infections) can prevent the induction of or break established tolerance.^14-19^ Notably, ischemia-reperfusion injury (IRI) is a critical determinant for an inflammatory reaction and initiation of rejection of the graft tissue after transplantation.^20,21^ Similarly to infections, extended graft ischemia is associated with reduced efficacy of CTLA4-Ig in controlling rejection.^22,23^

We hypothesized that multiple inflammatory cytokines provide alternative costimulatory signals (Signal 3) that compensate the inhibition of conventional costimulation. As a corollary, simultaneous CD28 blockade and inhibition of inflammatory cytokines would enhance costimulation blockade to promote lasting acceptance of a transplant. We therefore focused on the inhibition of cytokine signaling to contemporaneously block families of inflammatory cytokines with one drug (more cost effective than blocking multiple cytokines). Our objective was to delineate the synergism between CD28 blockade and inhibition of the JAK/STAT signaling pathway via the Janus kinase (JAK) 1 and 3 inhibitor Tofacitinib (Tofa).^24-27^ Tofa prevented allograft rejection in non-human primates, and in multiple randomized trials of human renal transplantation, it was as effective as tacrolimus.^28-30^ However, the safety profile of the prolonged use of Tofa resulted poorer, promoting discontinuation of its clinical investigation in transplantation. Rather than being used as chronically administered drug, we reasoned that Tofa could be used in short-term applications to potentiate the therapeutic efficacy of CTLA4-Ig and maintain an excellent safety profile.

Our results indicate that indeed Tofa unleashes the full immunomodulatory potential of CTLA4-Ig in inflammatory settings. This effect is achieved through effects on both APC and T cells, sustaining the impact of blocking Signal 2 by CTLA4-Ig. In our model of murine heterotopic heart transplantation, a short course administration of Tofa and CTLA4-Ig promoted long term allograft survival. More importantly, such a treatment exerted transplant survival even when the donor heart was kept ischemic for a prolonged time. Unexpectedly, the combination of Tofa and CTLA4-Ig promoted the accumulation and activity of regulatory T cells (Tregs). Overall, our results suggest that Tofa should be considered as an alternative (or complementary) to CD40 pathway blockade for maximization of the efficacy of CTLA4-Ig and safer management of transplant rejection.

## 2. Materials & Methods

### 2.1 Mice

BALB/c (H2-K^d^) and C57BL/6 (B6, H2-K^b^) mice were purchased from Charles River (Frederick, MD). Animals were housed in specific pathogen-free conditions and experiments conducted in accordance with National Institutes of Health guidelines, and with approval by the JHU Animal Care and Use Committee.

### 2.2 Reagents

Complete medium (CM) consisted of RPMI-1640 (Invitrogen, Carlsbad, CA) supplemented as previously described.^31^ CTLA4-Ig (Abatacept) was purchased from Johns Hopkins University Pharmacy. Tofacitinib and Ruxolitinib were purchased from LC Laboratories (Woburn, MA).

### 2.3 Generation of bone marrow derived dendritic cells and MATSup

Mouse bone marrow dendritic cells (DC) were generated as previously described.^32^ CD11c+ DC were purified by magnetic separation (EasySep Positive Selection; Stemcell Technologies, Cambridge, MA). Mature-DC were induced by adding LPS (200 ng/ml; E. coli K12; Invivogen, San Diego, CA) in the last 18h of culture. The supernatant from this LPS-DC culture was also collected and used as a source of inflammatory cytokines (**MATSup**) in T-cell proliferation assays. Fixed DCs were generated by exposure to a 2%-PFA solution for 10 min.

### 2.4 T cell isolation

Spleen and lymph nodes were harvested, and total T cells isolated via magnetic-bead negative selection as previously described.^31^ For suppression assays, Treg (CD4+CD25+) were isolated

from CD4-T cells following the protocol described in the EasySep PE-selection kit (STEMCELL technologies). Cells from the negative fraction (CD4+CD25-) were used as effectors.

### 2.5 *In vitro* Activation and Suppression Assays

For proliferation assays, freshly isolated T cells were stained with either CFSE or Violet trace (Thermo Fisher Scientific) following manufacturer’s instructions, and cocultured for 3-4 days with: allogenic DC; autologous DC and soluble αCD3 (0.05 μg/ml); αCD3/28-coated dynabeads (1:1 cell:bead ratio) or plate-bound αCD3/28 antibodies (BD Biosciences), and the indicated concentrations of Tofacitinib, CTLA4-Ig, and MATSup. For *in vitro* T cell-activation assay, T cells were activated for 24h and the percentage of IL-2-secreting cells detected by flow cytometry (IL-2-PE cytokine secretion assay, Miltenyi). For Suppression Assays, CFSE-stained T cells were stimulated in the presence of Tregs with the addition of Tofacitinib or Ruxolitinib. The trace-dye dilution profile was interpolated to calculate the precursor frequency using FlowJo proliferation analysis tool. The precursor frequency mean was normalized to the no-addition group.

### 2.6 Flow cytometry staining

Cells were stained with a fixable life/dead probe (Thermo Fisher Scientific) to exclude nonviable populations. In proliferation assays, cells were stained with anti-CD4 (GK1.5) and anti-CD8 (53-6.7) (Biolegend). For detection of intracellular cytokines, PMA+ionomycin (Cell-stimulating cocktail), and Brefeldin A (Protein Transport inhibitor cocktail; all from Thermo Fisher Scientific) were added the last 4h of culture. After surface staining, cells were fixed, permeabilized, and stained with antibodies anti-Foxp3 (FJK-16s), IFN-γ (XMG1.2) and TNF-α (MP6-XT22) (Thermo Fisher Scientific). In DC-maturation experiments, antibodies anti-CD11c (N418), anti-MHC-II (M5/114.15.2), anti-CD40 (1C10), anti-CD80 (16-10A1) and anti-CD86 (GL1) (Thermo Fisher Scientific) were used. Samples were acquired on an LSR-II cytometer (BD Biosciences), and results analyzed using FlowJo-X software (FLOWJO LLC, Ashland, OR).

### 2.7 Heterotopic heart transplantation

Intra-abdominal heart transplantation was performed from B6 to BALB/c mice as previously published.^33^ Graft was monitored by daily palpation and rejection defined as loss of heartbeat. Where indicated, mice received 500 μg CTLA4-Ig on post-operative-days POD0, 2, 4 and 6 and 15 mg/kg Tofacitinib via oral gavage twice-daily from POD0 to 6. Clinically relevant cold ischemia was recapitulated by incubating the allograft in ice-cold saline for 4h.

### 2.8 Isolation of graft-infiltrating cells

Explanted hearts were cut into pieces and digested for 30 min at 37C with 1 mg/ml of collagenase IV (Worthington) and 0.01% DNase I (Roche). Non-parenchymal cells were enriched via density interphase with 23.5% histodenz-PBS solution (MilliporeSigma) and an equal volume of CM. After centrifugation, the buffy-coat was collected, and characterized by flow cytometry.

### 2.9 Histology

Hearts were cut longitudinally, fixed in 10% formalin, and embedded in paraffin. Sections were stained with H&E. Slides were visualized on a Zeiss Axioplan-2 microscope and scanned with ProgRes CapturePro-2.7.6 software (JENOPTIK, MI). Lesions were graded by a blinded pathologist, based on the percentage of infiltration observed in the interstitial tissue as previously reported.^34^

### 2.10 Statistical analysis

Differences in transplant survival were assessed using a Mantel-Cox log-rank test. Statistical significance in *in vitro* assays was calculated using either unpaired Student’s *t* test or one-way ANOVA followed by Tukey post-test. Results are expressed as means ± SEM and analysis were performed with Graph Prism 7 (GraphPad, San Diego, CA). A *p*-value <0.05 was considered statistically significant.

## 3. Results

### 3.1 Pro-inflammatory cytokines promote T cell activation and counter the anti-proliferative effect of CTLA4-Ig

To test the hypothesis pro-inflammatory cytokines can provide an alternative costimulatory signal and counter the regulatory effect of CTLA4-Ig, we implemented an *in vitro* CFSE-T cells activation assay.^32^ This system demonstrated an expected dose-dependent reduction in T cell proliferation in presence of CTLA4-Ig (**Figure 1A and Supplementary Figure 1**),^9^ and the extent of inhibition was comparable between CD4 and CD8 T cells. To simulate an inflammatory environment, we added the supernatant of maturing DC (MATSup; see Materials and Methods). The addition of MATSup neutralized completely the inhibitory effect of CTLA4-Ig on both CD4 and CD8 T cells (**Figure 1A and Supplementary Figure 1**).

**Figure 1.**
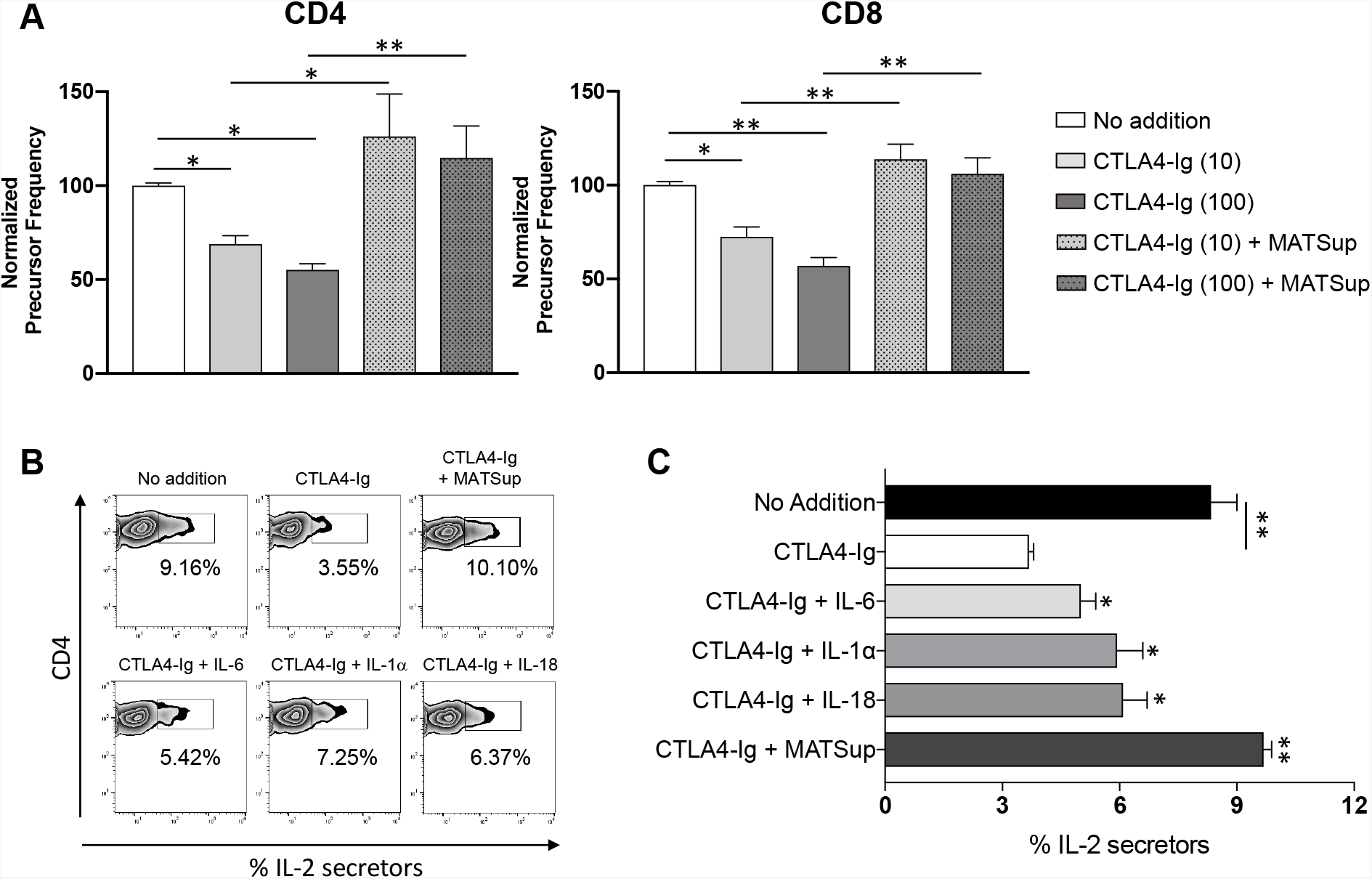
Inflammatory cytokines provide signals to T cells that counter CTLA4-Ig inhibition. **(A)** CFSE-labeled B6 T cells were stimulated for 72 h with soluble αCD3 and syngeneic DCs in presence of CTLA4-Ig (10 or 100 μg/ml) and MATSup. The extent of proliferation was extrapolated from the CFSE dilution profile, measured by flow cytometry, and compared among the different groups. (**B)** Representative dot-plots of IL-2 secretion (capture assay) at 24h of in vitro stimulation with soluble αCD3 and syngeneic DCs and **(C)** cumulative results obtained when CTLA4-Ig plus the indicated cytokines (IL-6, 20 ng/ml; IL-18, 10 ng/ml; IL-1α, 10 ng/ml) or MATSup, were added to the culture. Data shown in (A, C) are averages from n=4 independent experiments and expressed as precursor frequency normalized to the untreated condition (A) or as percentage of CD4 T secreting cells compared to the CTLA4-Ig only treated (C) ± SEM, *\*p* <0.05 and *\*\*p* <0.01 one-way analysis of variance (ANOVA) followed by Tukey post-test.

The effect of MATSup could be justified by the ability of pro-inflammatory cytokines to enhance the proliferation of T cells escaping inhibition by CTLA4-Ig, or by the hypothesized capacity to promote costimulation-independent activation. To distinguish between these two scenarios, we employed an “IL-2 capture assay” to determine the percentage of IL-2-secreting T cells present after 24h of stimulation – an early indicator of T cell activation, independent of cell proliferation.^35^ We chose three cytokines known to be abundant in the supernatant of maturing DC: IL-6, IL-1α, and IL-18. As shown in **Figure 1B-C**, CTLA4-Ig significantly reduced the percentage of IL-2 producing T cells. However, each cytokine investigated limited the inhibitory effect of CTLA4-Ig. More importantly, the presence of MATSup enabled a complete recovery of T cell activation, sustaining the hypothesis that pro-inflammatory cytokines promote CD28-independent activation of T cells.

### 3.2 Costimulation-independent activation of T cells depends on JAK-STAT signaling

Our results suggested the need for targeting multiple molecules to preserve the suppressive activity of CTLA4-Ig in an inflamed environment. We then repeated the CFSE-T cell proliferation assay in the presence of CTLA4-Ig+/-MATSup, with or without the addition of the JAK3/1 inhibitor Tofacitinib (Tofa).^24,27^ In the absence of inflammatory cytokines, both Tofa and CTLA4-Ig inhibited CD4 and CD8 proliferation (**Figure 2 and Supplementary Figures 2 and 3**). Their combination had an additive effect, exerting an even more profound inhibition of proliferation. With the addition of MATSup, neither agent alone fully regulated T cell activation (**Figure 2 and Supplementary Figures 2 and 3**); though, Tofa still had a statistically significant inhibitory effect on CD8 T cells. Notably, the combination Tofa+CTLA4-Ig had a profound effect in the presence of inflammatory cytokines, restoring the same level of inhibition exerted by CTLA4-Ig in absence of MATSup, in both T cell subpopulations. These results indicated that cytokines signaling through the JAK/STAT pathway has a key role in the impairment of CTLA4-Ig suppression.

**Figure 2.**
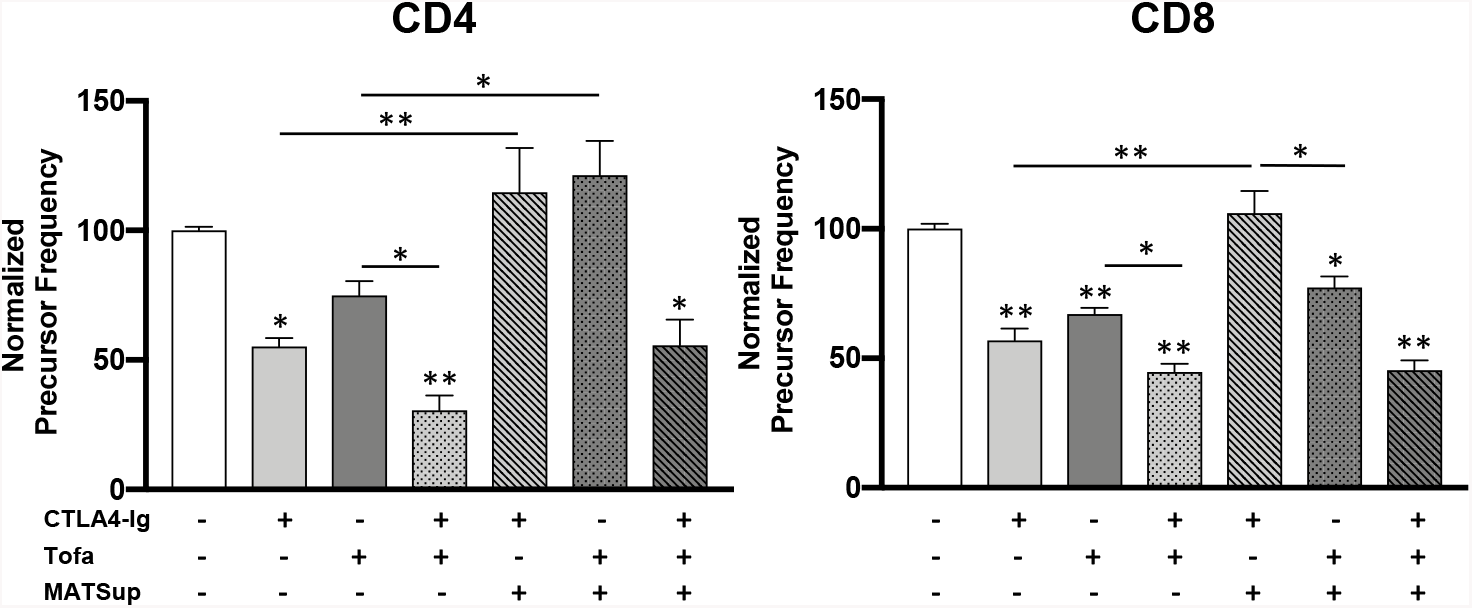
The JAK-STAT signaling pathway has a major role in co-stimulation independent T cell activation. CFSE-labeled T cells purified from B6 mice were stimulated for 72 h with soluble αCD3 (0.05 μg/ml) and syngeneic DCs with the addition, where indicated, of CTLA4-Ig (100 μg/ml), Tofa (1 μM) and MATSup. Proliferation was then measured by flow cytometry. Data shown is averaged from n=4 independent experiments and expressed as precursor frequency normalized and compared to the untreated condition or between samples (where indicated with connecting line) ± SEM, *\*p* <0.05 and *\*\*p* <0.01 one-way analysis of variance (ANOVA) followed by Tukey post-test.

### 3.3 Tofa impacts DCs function

The observed modulation of T cell proliferation by Tofa could derive from an impact on either T cells or DC (or both). Proinflammatory cytokines reportedly contribute to the activation and maturation of DCs.^17,18,36^ As suggested for the general class of JAK inhibitors,^37^ Tofa could interfere with the DC maturation process, consequently affecting their stimulatory capacity. To test this, we induced DC maturation with LPS in the presence or absence of Tofa. We then quantified the surface expression of key maturation markers and the release of inflammatory cytokines (**Figure 3A-B**). LPS+Tofa conditioned-DC demonstrated decreased expression of CD40, CD80, and CD86 compared to LPS matured-DC whereas, interestingly, both groups equally upregulated MHC-II **(Figure 3A)**. Correlatively, the exposure of DC to Tofa reduced the accumulation of IL-6, IL-1α, and TNF-α on a dose-dependent fashion **(Figure 3B)**.

**Figure 3.**
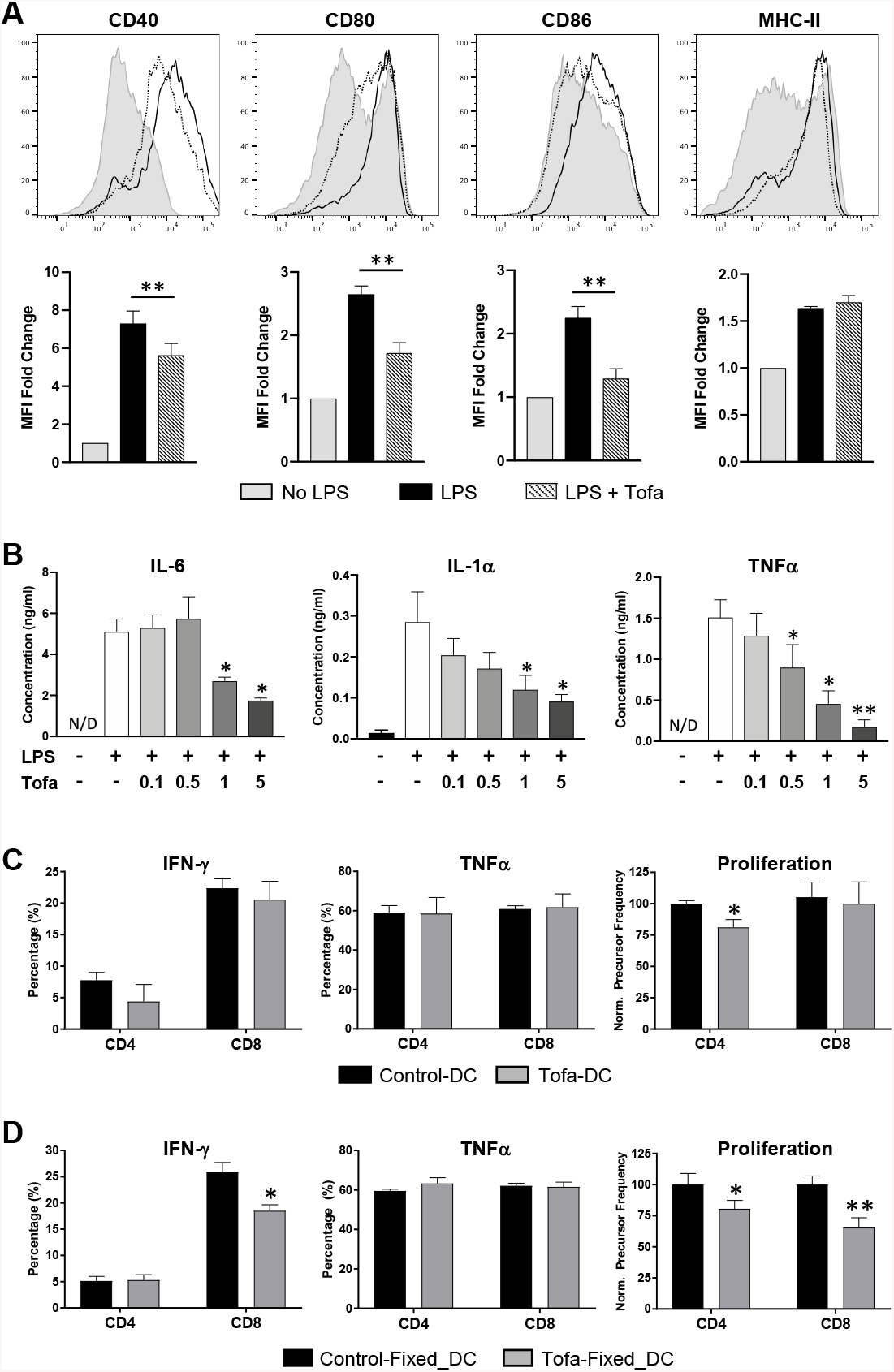
JAK inhibition impacts DC maturation. **(A-C)** Bone marrow derived B6 DC were either left untreated or exposed overnight to LPS +/- Tofa. **(A-top)** Representative histograms showing surface expression of the maturation markers CD40, CD80, CD86, and MHC-II determined by flow cytometry in CD11c+ cells. **(A-bottom)** Graph shows combined results from n=6 independent experiments expressed as the MFI fold change normalized to the untreated condition ± SEM, *\*p* <0.05 and *\*\*p* <0.01 unpaired Student’s *t-*test. (**B**) Supernatants from DC matured overnight with LPS in presence of a titration of Tofa (0.1-5 μM) were tested for accumulation of the indicated cytokines (IL-6, IL-1α, TNF-α) by ELISA. Results are representative of n=3 independent experiments and expressed as average concentration in the supernatant (in ng/ml) ± SEM, *\*p* <0.05, two-tailed unpaired Student’s *t-*test. **(C)** CFSE-labeled B6 T cells were cultured for 4 days with allogeneic BALB/c-DC (1:20, DC:Tcell ratio) that were previously matured with LPS in the presence or absence of Tofa and then washed before co-culture. **(D)** CFSE-labeled B6 T cells were cultured for 4 days with allogeneic BALB/c-DC (1:20, DC:Tcell ratio) that were previously matured with LPS in the presence or absence of Tofa, and then fixed with 2%-PFA before co-culture. (**C-D**) Induced proliferation and intracellular cytokine levels (post PMA/Ionomycin re-stimulation) were measured by flow cytometry. Data shown is an average of n=3 independent experiments and expressed as the precursor frequency value normalized to the proliferation induced in the LPS-DC cultured group, or as the percentage of cytokine secreting CD4/CD8 T cells ± SEM with *\*p* <0.05, *\*\*p* <0.01 two-tailed unpaired Student’s *t-*test.

We then tested the impact of this altered DC maturation state on T cell activation. BALB/c-DC matured with LPS+/-Tofa were washed and used to stimulate B6-T cells in a mixed leucocyte reaction (MLR) (**Figure 3C**). Surprisingly, we measured a limited decrease in alloreactive CD4-T cell proliferation when cells were stimulated by Tofa-conditioned LPS-matured DC; CD8 T cell were not affected at all. Similarly, no changes were observed in the production of IFN-γ or TNF-α by either subset (**Figure 3C**). We argued that these results could be justified by a reversibility of the inhibition of DC maturation by Tofa (present only during DC conditioning). To test this hypothesis, LPS+Tofa conditioned-DC were rested in drug-free media for additional 24h. Analysis of CD40L, CD80 and CD86 expression indicated a full recovery of the maturation status **(Supplementary Figure 4A)**. We then repeated the MLR using DC fixed with PFA after conditioning, so to fix the effect of Tofa (**Supplementary Figure 4B**). In these conditions, fixed-Tofa-conditioned LPS-matured DC induced a decreased proliferation in both CD4 and CD8 T cells and IFN-γ secretion by the latter (**Figure 3D**). These results indicated that Tofa has a profound impact on DC maturation, which affects T cell activation, and its effect is rapidly reversible.

As Tofa prevents the release of proinflammatory mediators by DC, we tested if this altered secretion pattern would have an impact on T cell activation. We collected the supernatants of DC exposed to LPS with or without Tofa (MATSup^Tofa^ and MATSup) and assessed their effect on T cell proliferation in a DC-free *in vitro* activation system (**Figure 4A**). In concordance with the reduced content of pro-inflammatory cytokines, MATSup^Tofa^ induced a significantly lower increase in T cell proliferation than conventional MATSup. However, when we accounted for the possible contribution of Tofa carried over in the MATSup^Tofa^ (by adding an estimated equivalent amount to the MATSup – MATSup+Tofa), the difference was still statistically different in CD4-but not in CD8-T cell proliferation (**Figure 4A**).

**Figure 4.**
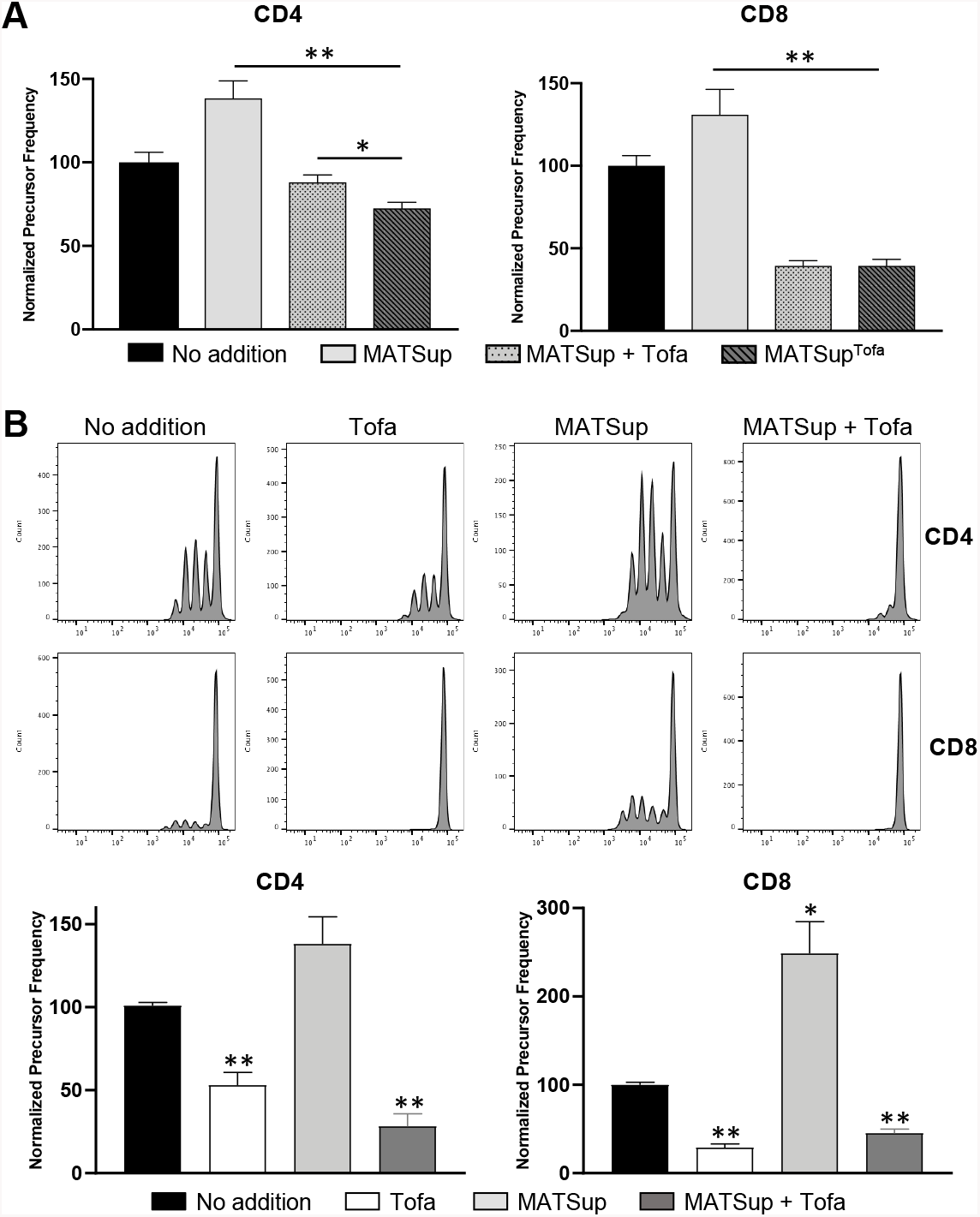
Tofa directly inhibits T cell proliferation. **(A)** CFSE-labeled purified B6 T cells were cultured for 72h with αCD3/αCD28 dynabeads (1:1 cell:bead ratio) with the addition of the supernatant from: DC exposed to LPS (MATSup), DC exposed to LPS in presence of Tofa (MATSup^Tofa^), or DC exposed to LPS with delayed addition of Tofa to the media (MATSup + Tofa). Proliferation was then measured by flow cytometry. Data shown is averaged from n=3 independent experiments and expressed as the average precursor frequency in CD4 or CD8 normalized to the no-addition condition ± SEM, *\*p* <0.05 and *\*\*p* <0.01 two-tailed unpaired Student’s *t-*test. **(B)** CFSE-labeled purified B6 T cells were stimulated for 72h with plate coated αCD3 (2.5 μg/ml) and soluble αCD28 (0.01 μg/ml) in the presence of Tofa (1 μM) and/or MATSup. Proliferation was then measured by flow cytometry. **(B-top)** Representative histograms showing proliferation by cell-trace dye dilution. **(B-bottom)** Cumulative results from n=4 independent experiments with data expressed as the average precursor frequency in CD4 or CD8 populations normalized to the no-additions condition ± SEM, *\*p* <0.05 and *\*\*p* <0.01 two-tailed unpaired Student’s *t-*test.

### 3.4 Tofa directly inhibits T cell activation

We then tested if Tofa directly impacts T cell activation. We stimulated T cell with plate-bound αCD3 and suboptimal concentration of αCD28 (to simulate the effect of the addition of CTLA4-Ig). In these settings, the presence of MATSup increased T cell proliferation and countered the suboptimal level of αCD28 (similarly to the presence of CTLA4-Ig in Figure 1A and 2) (**Figure 4B**). Strikingly, Tofa showed a profound suppressive effect, inhibiting both CD4 and CD8 T cells proliferation (with a greater impact on the latter). Notably, this inhibition was maintained in presence of MATSup. These results demonstrate that Tofa has a direct inhibitory effect on T cells that counters the costimulation-independent activation promoted by inflammatory cytokines.

### 3.5 Tofa synergizes with CTLA4-Ig in promoting cardiac transplant survival and extends its efficacy to ischemic grafts

Supported by the *in vitro* data, we assessed if a short course of Tofa could synergize with CTLA4-Ig to promote transplant survival in a full-mismatch murine heterotopic heart transplantation model.^38^ Our regimen consisted of CTLA4-Ig administration on POD0, 2, 4, 6 and Tofa twice daily for the period POD0-6. Notably, the CTLA4-Ig+Tofa combination significantly extended allograft survival compared to CTLA4-Ig alone (MST=158 vs 36 respectively - **Figure 5A**).

**Figure 5.**
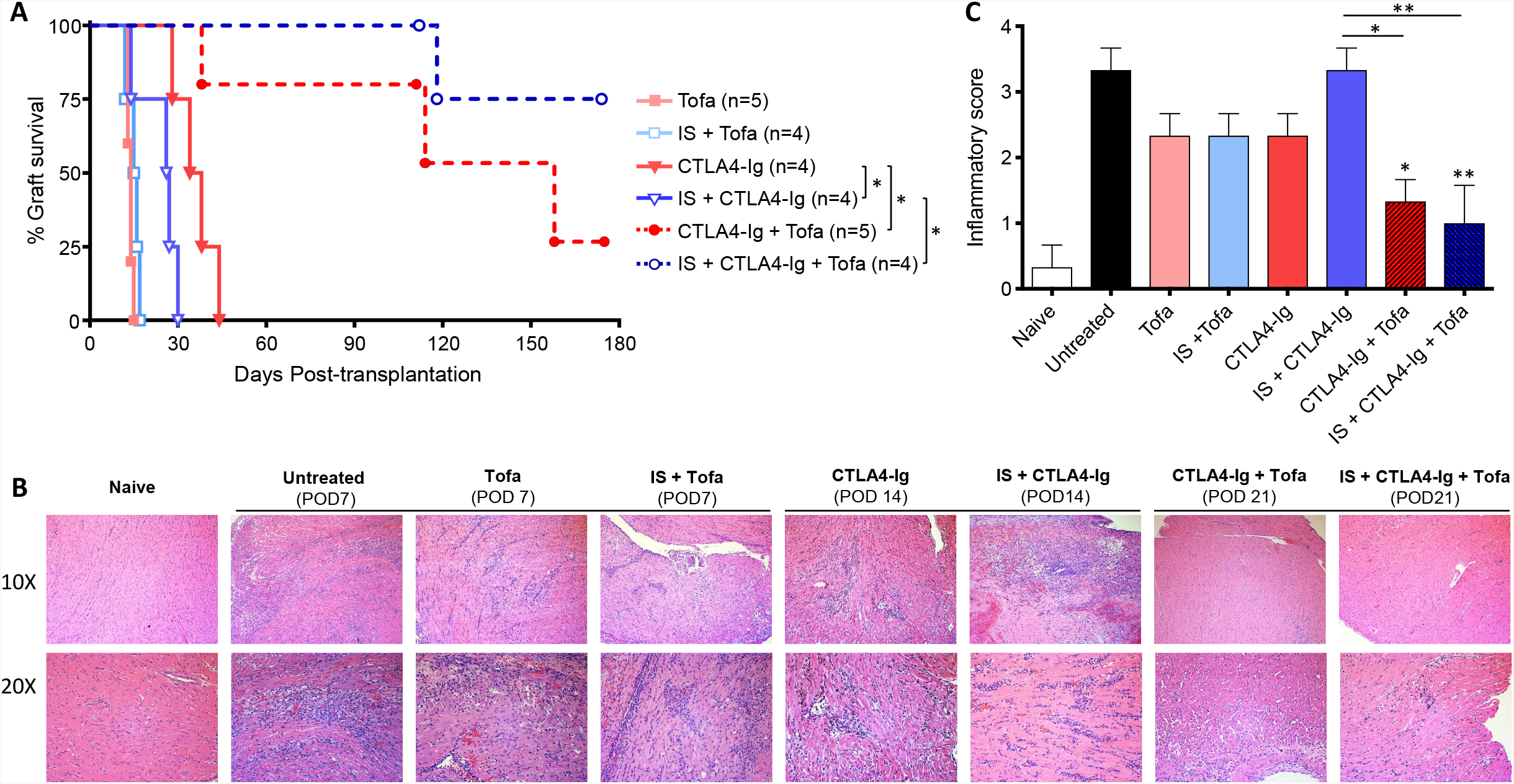
The combined use of Tofa and CTLA4-Ig promotes long-term transplant survival and equally protects ischemic grafts. B6 hearts were either transplanted immediately into BALB/c mice or first subjected to a 4h cold ischemia (IS) before grafting. Recipients were then treated with 500 μg CTLA4-Ig on days 0, 2, 4 and 6 post-transplantation, alone or in combination with 15 mg/kg Tofa (Tofa-citrate, diluted in 0.5% methylcellulose/0.025% Tween-20) via oral gavage twice-daily from day 0 to 6. **(A)** Allograft survival curves in the different treatment groups (n=4-5 animals each). Differences are expressed as *\*p* <0.05, Log-Rank test. **(B)** Representative H&E histology samples showing leucocytic infiltration prior to transplant rejection – Untreated, Tofa and IS+Tofa (POD 7); CTLA4-Ig and IS-CTLA4-Ig (POD14); and CTLA4-Ig+Tofa and IS+CTLA4-Ig+Tofa (POD 21). **(C)** Composite histological inflammatory scores (n=3 mice/group) including all the groups listed in B. Differences between groups are expressed as *\*p* <0.05 and **p<0.01 one-way analysis of variance (ANOVA) with Tukey post-test.

Histological examination indicated that while CTLA4-Ig-only group presented multifocal moderate lymphohistiocytic myocarditis and endocarditis with perivascular inflammation, allografts receiving CTLA4-Ig+Tofa had a milder affectation, though not statistically significant (**Figure 5B-C**). To further investigate the clinical applicability of the combined regimen, we tested its efficacy when using ischemic hearts, a condition characterized by an increased accumulation of proinflammatory cytokines in the tissue.^22,23^ In these settings, CTLA4-Ig monotherapy showed a reduced protective effect compared to no-ischemia and this result was confirmed in our strain combination (MST=26.5 vs 36 **Figure 5A**). Remarkably, the administration of CTLA4-Ig+Tofa prolonged the survival of ischemic hearts to the same extent as non-ischemic ones (MST>175 vs MST=158 - **Figure 5A**). Concordantly, histology samples showed a moderate lymphohistiocytic inflammation in the CTLA4-Ig-only group vs only a focal mild infiltrate in the group receiving the full protocol (**Figure 5B-C**). Administration of Tofa alone minimally extended transplant survival, in both ischemic and non-ischemic conditions (MST=15.5 and 14 respectively - **Figure 5A**). These results suggest that a short treatment with Tofa sustains the therapeutic efficacy of CTLA4-Ig.

### 3.6 Tofa+CTLA4-Ig promote the intra graft accumulation of regulatory T cells and reduce the differentiation of TH1 effectors

To understand the mechanism behind the therapeutic effect of the combined regimen, we analyzed the composition of allograft infiltrating T cells. In line with the survival data, CTLA4-Ig promoted the accumulation of Treg (**Figure 6A**). The combination of Tofa and CTLA4-Ig promoted an even higher Treg increase. Interestingly, in both CTLA4-Ig and Tofa+CTLA4-Ig treatment groups receiving an ischemic graft, the proportion of Foxp3+ T cells was comparable. We also assessed T_H_1 differentiation in the spleen by measuring the percentage of IFN-γ producing CD4-T cells. As shown in **Figure 6B**, CTLA4-Ig-only treatment had a moderate impact on the accumulation of TH1 cells, while the addition of Tofa prevented completely their formation. This effect was retained even in the recipients of ischemic hearts. These results suggest that Tofa+CTLA4-Ig promotes a robust regulation of the formation of alloreactive effectors while sustaining the accumulation of intra-graft Treg that control the rejection response.

**Figure 6.**
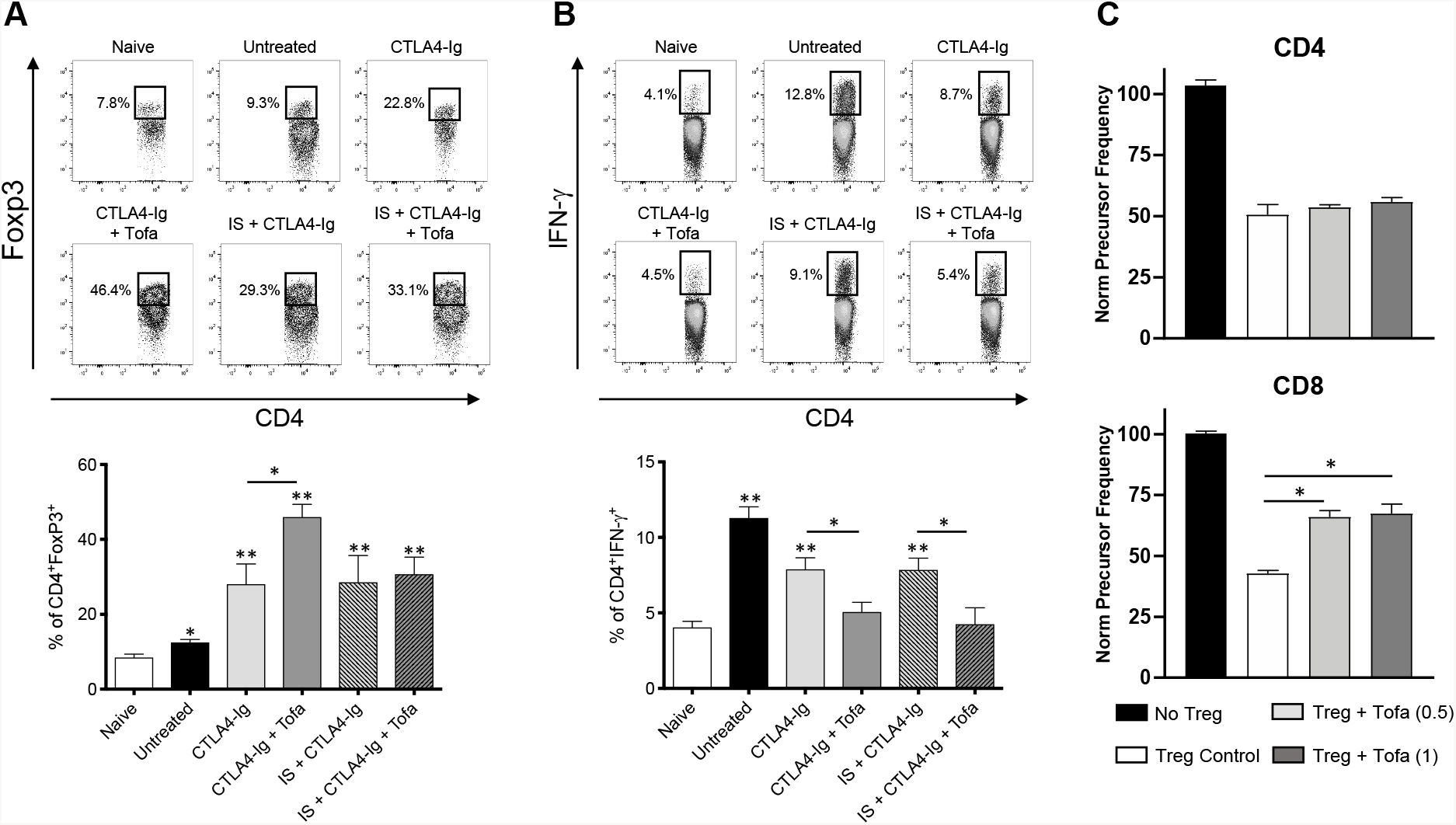
CTLA4-Ig and JAK inhibition promote intra-graft Tregs accumulation and decrease the differentiation of TH1 cells. **(A**,**B)** Heart allografts and spleens were harvested from recipient animals (subjected to the same treatments indicated in Figure 5) prior to transplant rejection; Untreated (POD 7); CTLA4-Ig and CTLA4-Ig+Tofa (POD 21); IS+CTLA4-Ig and IS+CTLA4-Ig+Tofa (POD 14), and percentages of intra-graft Tregs and splenic IFN-γ+ CD4 T cells (post PMA/Ionomycin re-stimulation) were characterized by flow cytometry. **(A)** Representative data and cumulative results showing the percentage of allograft infiltrating CD4+Foxp3+ cells. **(B)** Representative data and cumulative results of the percentage of splenic CD4+IFN-γ+ cells. Data shown in (A, B) are the averages from n=3 independent experiments each generated from 3 individual mice/group and are expressed as percentage ± SEM, *\*p* <0.05 and *\*\*p* <0.01 two-tailed unpaired Student’s *t-*test. **(C)** Treg suppression assay. CFSE-labeled CD4^+^CD25^-^ T cells were stimulated with soluble αCD3 plus syngeneic DC and cocultured with CD4^+^C25^+^ Tregs (2:1 ratio Teff:Treg), in presence of the indicated concentrations of Tofa (0.5-1 μM). The suppression of proliferation induced by Tregs was measured via flow cytometry. Data shown is averaged from n=3 independent experiments each generated from individual mice, and expressed as precursor frequency normalized to the proliferation measured without Treg ± SEM, *\*p* <0.05, two-tailed unpaired Student’s *t-*test.

Because of the observed accumulation of Treg, we assessed if Tofa was permissive of Treg-activity by employing an *in vitro* CFSE-based Treg suppression assay.^39^ The resulting data (**Figure 6C**) demonstrated that Tofa does not interfere with Treg suppression of CD4-T cell proliferation, but it partially reduces the regulation of CD8-T cells (**Figure 6C**). To further confirm that Treg activity is only minimally affected by JAK inhibition, we repeated the same suppression assay in presence of Ruxolitinib, a JAK1/2 inhibitor recently approved by the FDA for the treatment of steroid-refractory acute graft-versus-host-disease.^40^ Ruxolitinib did not affect the capacity of Treg to regulate CD4 or CD8 T cells (**Supplementary Figure 5**), indicating that the combination of JAK-inhibition with CTLA4-Ig is permissive of Treg activity.

## 4. Discussion

Our data further rationalize the observations that inflammatory cytokines impede costimulation blockade regimens^15,16,41^ and suggest that the addition of short-courses of Tofa should be considered for maximizing the therapeutic efficacy of Belatacept.^10,42^

T cells require a third signal, along with TCR and costimulatory receptors engagement, to build a productive response.^43-45^ We report here that multiple proinflammatory cytokines (IL-6, IL-18 and IL-1α) have the ability to compensate for reduced costimulatory signaling. When present in combination (MATsup), pro-inflammatory cytokines act in synergy and completely override the inhibition of T cell proliferation induced by CTLA4-Ig.^9^ This outcome reinforces the understanding that Signal 2 (costimulation) and Signal 3 (cytokine signaling) can be complementary in T cell activation, and highlights the importance of blocking both signals to control alloreactive T cells.

Inflammatory cytokines use multiple signaling pathways to deliver their functions,^46^ and our results show that Tofa is very effective at suppressing a key pathway involved in T cell activation. As first generation JAK inhibitor, Tofa targets JAK3/1/2 and can block the signaling of almost any cytokine using the JAK/STAT pathway, depending on the concentration used.^27,47^ Importantly, our results indicate that exposure of DC to Tofa significantly reduced their production of IL-1 and TNF-α in response to maturative stimuli. The signaling of these cytokines would not be inhibited by Tofa and we believe the prevention of their accumulation contributes indirectly to controlling T cell activation. Ultimately, our data show that Tofa fully restores the capacity of CTLA4-Ig to control T cell proliferation under inflammatory conditions, suggesting that inflammatory cytokines counteract CTLA4-Ig-mediated inhibition of T cell activation with a JAK/STAT-dependent mechanism. This is supported by our *in vivo* results, where the combined therapy extended the transplant survival achieved with CTLA4-Ig monotherapy, even in the group subjected to prolonged cold ischemia. It should be investigated if this effect extends to TLR-mediated blockade of tolerance previously reported.^18,23,48^

Functionally, Tofa affects both innate and adaptive immune responses by limiting the maturation of innate cells and inhibiting pathogenic TH1, TH2, and TH17 cells differentiation,27,49 properties that supported its use for autoimmune disorders (it is FDA approved for treatment of rheumatoid arthritis and ulcerative colitis) and transplantation.^24,26,27^ In concordance with previous publications^50,51^ we report that Tofa affects DC maturation, an effect probably mediated by its inhibition of JAK1/2 and STAT1/4, all involved in the response to LPS.^52-54^ Very interestingly, as also observed in human monocytes,^51^ Tofa exposure does not prevent MHC-II upregulation. The net result of such an altered phenotype would be the preservation of alloantigens presentation (Signal 1) with reduced Signal 2 and 3 (strengthening the effect of CTLA4-Ig) and ultimately translating into a tolerogenic DC phenotype.^55-59^ We believe these regulatory effects contribute to the extension of transplant survival we observed, and we are further investigating this mechanistic aspect.

Our *in vitro* results indicate a differential impact of Tofa on T cells activation when stimulated in presence of MATSup by αCD3+autologous DC vs αCD3/28. With DC, JAK inhibition reduced T cells proliferation of only a small degree (if any). With αCD3/28 stimulation, both CD4 and CD8 cells were susceptible to Tofa, with an almost complete loss of proliferation. A possible explanation is the different impact of JAK signaling in DC vs T cells. DC are not fully dependent on JAK signaling for maturation. Then, Tofa only partially affects their ability to provide Signal 2 and 3 to T cells, remaining able to provide enough costimulation for activation. Contrarily, T cells appear to be strongly dependent on the JAK/STAT pathway for activation when engaging exclusively CD3/CD28, making Tofa effective at inhibiting activation in these settings **(Figure 7)**.

**Figure 7.**
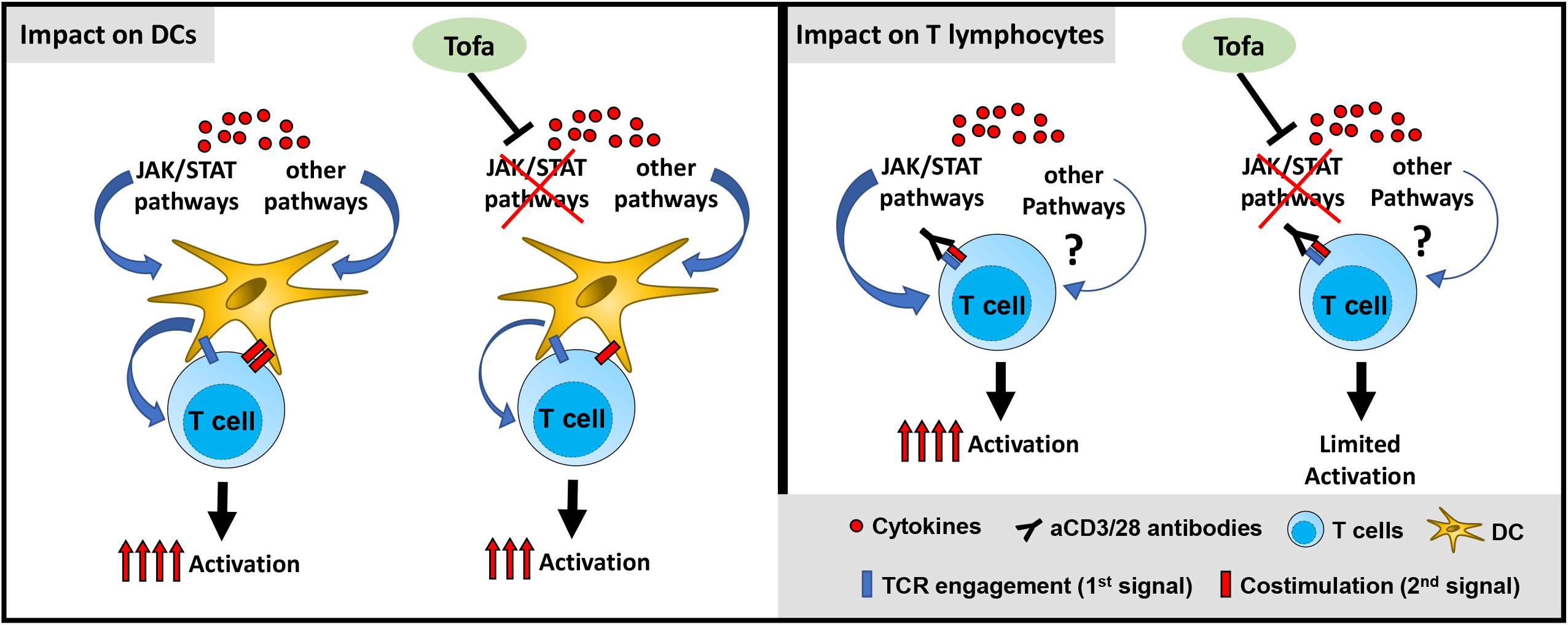
Proposed working model of Tofa effect on DCs and T cells.

In our transplant model, the combination of Tofa and CTLA4-Ig decreased the TH1-IFN-γ effector response, the population primarily responsible for acute rejection, even when the graft was subjected to prolonged ischemia.^60^ We believe this result derives from the capacity of Tofa to inhibit the activation of the TH1 lineage transcription factor T-bet,^27,28,61^ acting in synergy with CTLA4-Ig.^62^ We also report an increase in the proportion of intra-graft Treg, a condition associated with extended transplant survival in multiple models.^63,64^ Differently from the effect on TH1 cells, the increase of intra-graft Treg was not evident when implementing ischemic hearts, despite the comparable extension of transplant survival. This difference suggests a possible negative impact of inflammation on Treg suppressor activity^65^ and recruitment that, however, does not impact graft survival with our treatment strategy. We believe all these positive effects are associated to the short-term administration of Tofa, an approach never considered before. Long-term administration of this drug has been associated to a decrease on Treg abundance.^28^ We believe our combination therapy creates the conditions to sustain the activity of Treg. Our results align with more recent publications that show that Tofa spare Treg function,^66^ and suggest this property could extended to other JAK/STAT inhibitors such as Ruxolitinib.

As standalone immunosuppressant Tofa demonstrated great inhibitory capacity,^28,29^ but safety concerns deriving from its long-term administration paused its further clinical investigation in transplantation.^67-69^ We propose a different utilization of Tofa: short-term administration intervals that parallel the infusion of CTLA4-Ig. This regimen would be more cost-effective than the combination of CTLA4-Ig with biologics like complement inhibitors^23,70^ or blocking antibodies,^22,41^ while preserving similar efficacy. In addition, the reported rapid normalization of blood parameters after treatment termination,^71^ and the quick reversibility of inhibition of DC phenotype we reported, indicate that Tofa allows great flexibility in patient management.^67-69^ With further elucidation of the mechanisms underlying the synergism between Tofa and CTLA4-Ig and with optimization of the administration protocol, e.g. by implementing a localized delivery of Tofa (a strategy we are currently exploring),^72^ we believe there is great potential for designing safer and highly effective strategies for management of transplant recipients.

## Abbreviations

APC: Antigen presenting cells
B6: C57BL/6 mouse strain
CM: Complete media
CTLA4: Cytotoxic T-Lymphocyte Associated protein 4
DC: Bone marrow-derived dendritic cells
IRI: Ischemia Reperfusion Injury
JAK: Janus Kinase
LPS: Lipopolysaccharide
MATSup: Supernatant of LPS-matured DC
MST: Mean survival time
TCR: T cell receptor
Tofa: Tofacitinib

## Acknowledgments

This work was supported by NIH grant 1R21HL127355, United States Army Medical Research Acquisition Activity (USMRAA) grant W81XWH-18-1-0789, and internal support from the Dep. of Plastic & Reconstructive Surgery (all to G.R.). The authors thank Sonia Santiago and Samiya Soto (laboratory managers), Xiaoling Zhang and Dixie Hoyle (JHU Ross flow cytometry core) for excellent technical assistance; the Division of Research Animal Resources (RAR) at Johns Hopkins University for animal husbandry and care; and JHU Phenotypic Core Facility for processing histology samples of transplanted heart tissues. The authors also thank Dr. Sarah Poynton (Dep. of Molecular and Comparative Medicine) for invaluable assistance with manuscript editing.

## Data Availability Statement

The data that support the findings of this study are available from the corresponding author upon reasonable request.

## Supporting information statement

Additional supporting information may be found online in the Supporting Information section at the end of the article

