## Supplementary Figure for "A short course of Tofacitinib sustains the immunoregulatory effect of CTLA4-Ig in presence of inflammatory cytokines and promotes long-term survival of murine cardiac allografts"

### Supplementary information

**Supplementary Figure S1.**

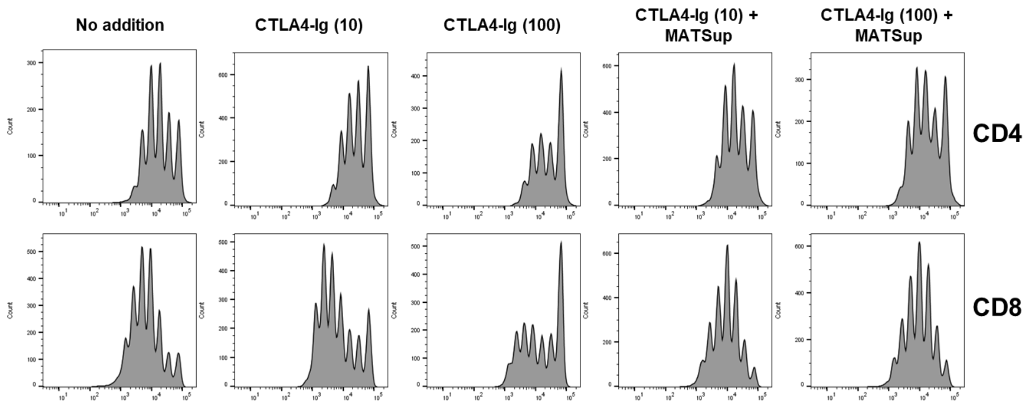


**Supplementary figure S1.- Representative histograms of proliferation in figure 1A.** Representative histograms showing the extent of T cell proliferation (CFSE-dilution) measured by flow cytometry in cultures where CFSE-labeled B6 T cells were stimulated for 72 h with soluble αCD3 and syngeneic DCs in presence of CTLA4-Ig (10 or 100 μg/ml) and MATSup.

**Supplementary Figure S2.**

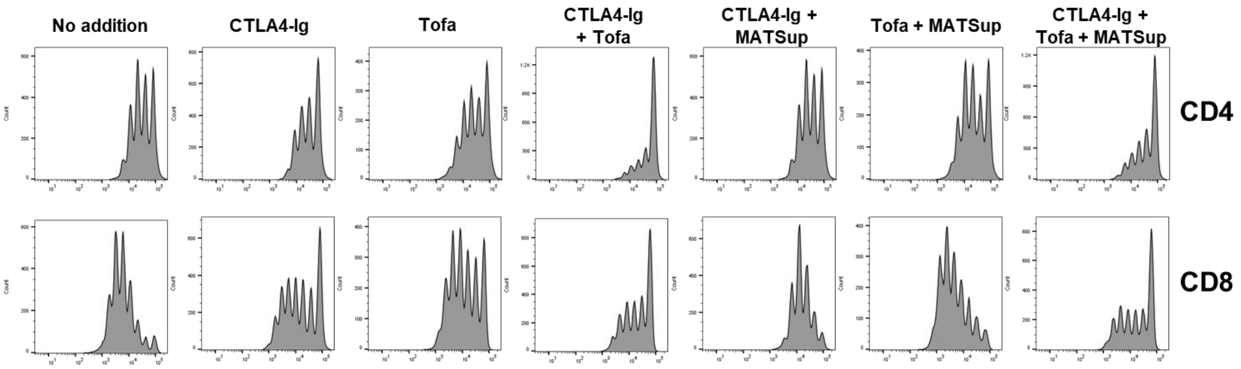


**Supplementary figure S2.- Representative histograms of proliferation in figure 2.** Representative histograms showing T cell proliferation (CFSE-dilution) measured by flow cytometry in cultures where CFSE-labeled B6 T cells were stimulated for 72 h with soluble αCD3 and syngeneic DCs with the addition, where indicated, of CTLA4-Ig (100 μg/ml), Tofacitinib (1 μM) and MATSup.

**Supplementary Figure S3.**

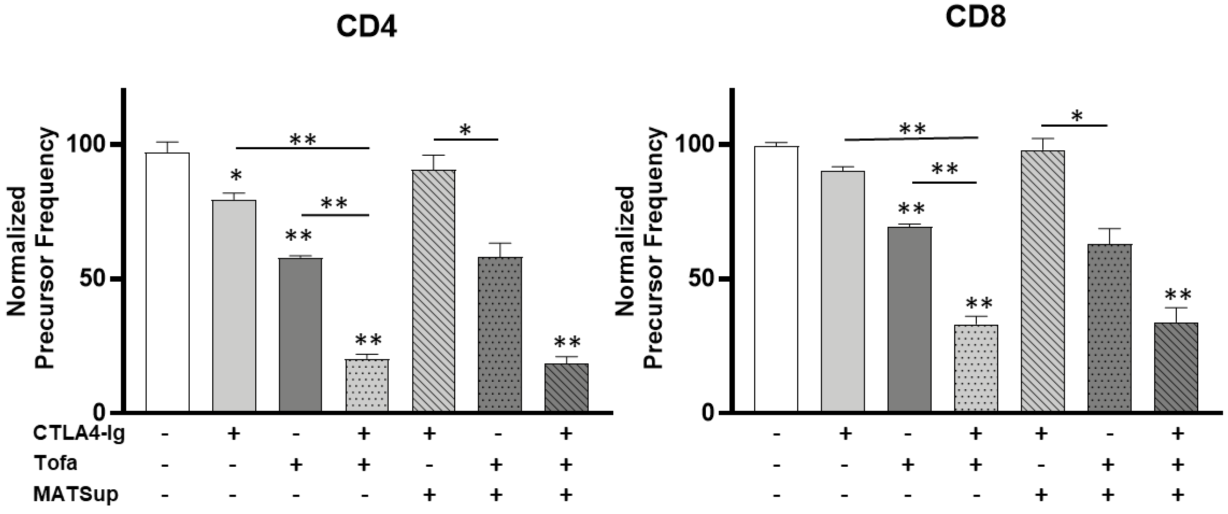


**Supplementary figure S3.- The JAK-STAT signaling pathway has a major role in co-stimulation independent activation of Balb/c T cells.** CFSE-labeled T cells purified from Balb/c mice were stimulated for 72 h with soluble αCD3 (0.05 μg/ml) and syngeneic DCs with the addition, where indicated, of CTLA4-Ig (100 μg/ml), Tofa (1 μM) and MATSup. Proliferation was then measured by flow cytometry. Data shown is averaged from n=3 independent experiments and expressed as precursor frequency normalized and compared to the untreated condition ± SEM, *$p$ <0.05 and **$p$ <0.01 one-way analysis of variance (ANOVA) followed by Tukey post-test.

**Supplementary Figure S4.**

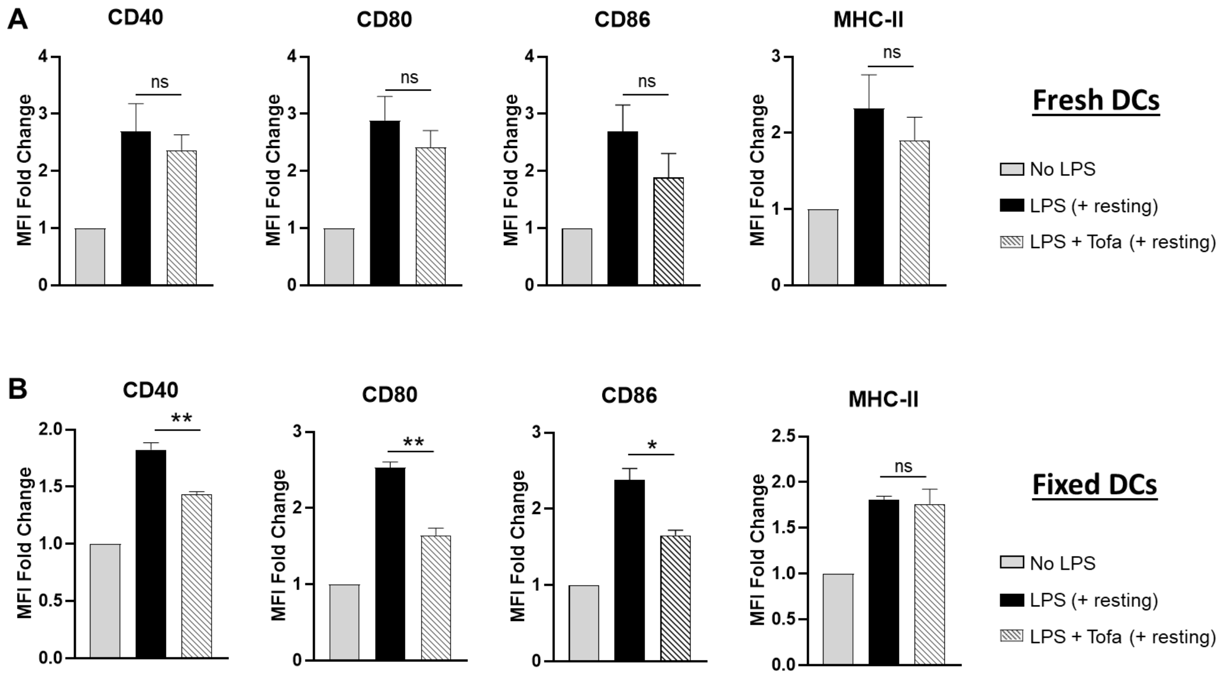


**Supplementary figure S4.- Reversibility of Tofa inhibition of DC maturation. (A)** B6 bone marrow derived DC were left untreated or exposed overnight to LPS +/- Tofa, washed, and then rested for additional 24h in media free of LPS or Tofa. **(B)** B6 bone marrow derived DC were left untreated or exposed overnight to LPS +/- Tofa, Fixed with 2% Paraformaldehyde (PFA), washed, and then rested for additional 24h in media free of LPS or Tofa. **(A, B)** Expression of the maturation markers CD40, CD80, CD86, and MHC-II after the resting phase was determined by flow cytometry in Live CD11c+ cells. Graph shows combined results from n=5 (A) or n=3 (B) independent experiments expressed as the average of MFI fold change normalized to the untreated condition ± SEM, *$p$ <0.05, two-tailed unpaired Student's $t$-test.

**Supplementary Figure S5.**

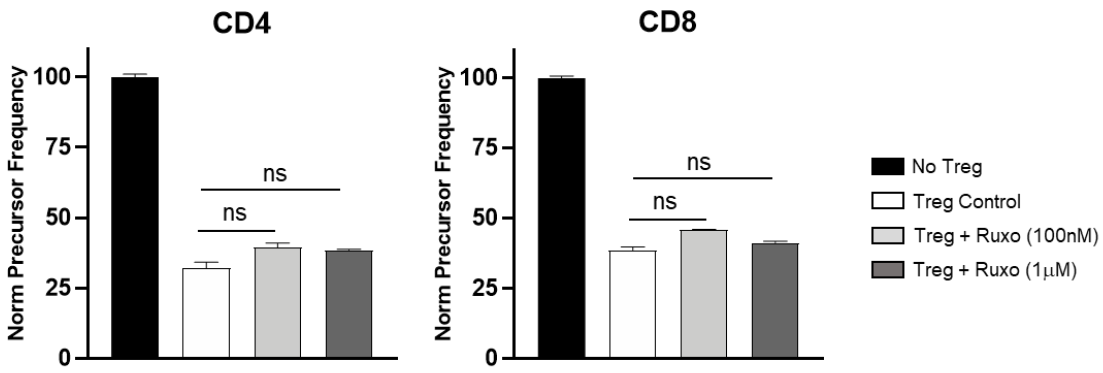


**Supplementary figure S5.- Ruxolitinib is permissive of Treg suppressive activity.** Treg suppression assay. CFSE-labeled $CD4^{+}CD25^{-}$ T cells were stimulated with soluble αCD3 and syngeneic DCs and cocultured with $CD4^{+}C25^{+}$ Tregs (2:1 ratio Teff:Treg), in the presence of different concentrations of Ruxolitinib (0.1-1 μM). The extent of suppression of proliferation induced by Tregs was measured by flow cytometry. Data shown are averaged from n=2 independent experiments and are expressed as precursor frequency normalized to the extent of proliferation measured in the no-Treg group ± SEM, *$p$ <0.05, *$p$ <0.01 two-tailed unpaired Student's *t*-test.
